# Emergence and Evolution of a Prevalent New SARS-CoV-2 Variant in the United States

**DOI:** 10.1101/2021.01.11.426287

**Authors:** Adrian A. Pater, Michael S. Bosmeny, Christopher L. Barkau, Katy N. Ovington, Ramadevi Chilamkurthy, Mansi Parasrampuria, Seth B. Eddington, Abadat O. Yinusa, Adam A. White, Paige E. Metz, Rourke J. Sylvain, Madison M. Hebert, Scott W. Benzinger, Koushik Sinha, Keith T. Gagnon

## Abstract

Genomic surveillance can lead to early identification of novel viral variants and inform pandemic response. Using this approach, we identified a new variant of the SARS-CoV-2 virus that emerged in the United States (U.S.). The earliest sequenced genomes of this variant, referred to as 20C-US, can be traced to Texas in late May of 2020. This variant circulated in the U.S. uncharacterized for months and rose to recent prevalence during the third pandemic wave. It initially acquired five novel, relatively unique non-synonymous mutations. 20C-US is continuing to acquire multiple new mutations, including three independently occurring spike protein mutations. Monitoring the ongoing evolution of 20C-US, as well as other novel emerging variants, will be essential for understanding SARS-CoV-2 host adaptation and predicting pandemic outcomes.

## Introduction

In early 2020 the World Health Organization declared that coronavirus disease 2019 (COVID-19), a potentially fatal respiratory infection caused by SARS-CoV-2, was a global pandemic (*1*). The high number of SARS-CoV-2 infections worldwide over time has presented the virus with ample opportunity to acquire new mutations. It has been suggested that some mutations already present a fitness advantage for the virus. Notably, the D614G mutation, observed early in the pandemic, is thought to increase the transmissibility of the virus (*2, 3*). The N501Y mutation of the spike protein (S) has been implicated in the rapid spread of new variants in the United Kingdom and South Africa (*4–6*). A growing number of spike protein mutations could enable immune evasion and reduced vaccine efficacy (*7*).

The U.S. has experienced a surge in cases during the third pandemic wave over the fall of 2020 and winter of 2020/2021. While many variables are likely to drive the increase in cases, it is possible that emergence of a more fit or transmissible SARS-CoV-2 variant could be a contributing factor (*5*). Restrictions in population movement during a global pandemic, as well as the rapid acquisition of multiple mutations, could drive emergence of novel region-specific variants. This evolutionary paradigm might explain the rise of distinct SARS-CoV-2 variants now being observed around the world during the COVID-19 pandemic (*5, 6, 8*).

Here we report the characterization of a SARS-CoV-2 variant, 20C-US, that emerged in and has remained mostly confined to the U.S. Its quiet rise to prominence among other circulating variants in the late summer and early fall of 2020 coincides with the third U.S. pandemic wave. Based on existing genomic data, we predict that this variant may already be the most dominant variant of SARS-CoV-2 in the U.S., likely accounting for the majority of COVID-19 cases. In addition to the five signature mutations of the 20C-US variant, new mutations continue to accrue. These include protease, nucleocapsid, and spike protein mutations that highlight the ongoing evolution of SARS-CoV-2.

## Results

### Genomic and phylogenetic and characterization of 20C-US, a prevalent new SARS-CoV-2 variant in the U.S

In response to anticipated genetic changes occurring in the SARS-CoV-2 virus, we began sequencing viral genomes for genomic epidemiology and surveillance. With sequencing focused on the U.S. upper Midwest in the state of Illinois, we generated full genome sequences from samples taken beginning in March 2020 to present. During phylogenetic reconstruction with our Illinois genome sequences, a particular branch within the 20C clade became noticeably more pronounced (**Fig. 1A**). We identified five closely co-occurring signature mutations that appeared synapomorphic to the new clade within 20C. These mutations resulted in amino acid changes of N1653D and R2613C in ORF1b, G172V in ORF3a, and P67S and P199L in the nucleocapsid (N) gene, the last of which also introduces a stop codon mutation at position Q46 of ORF14 (**Table 1**).

**Fig. 1.**
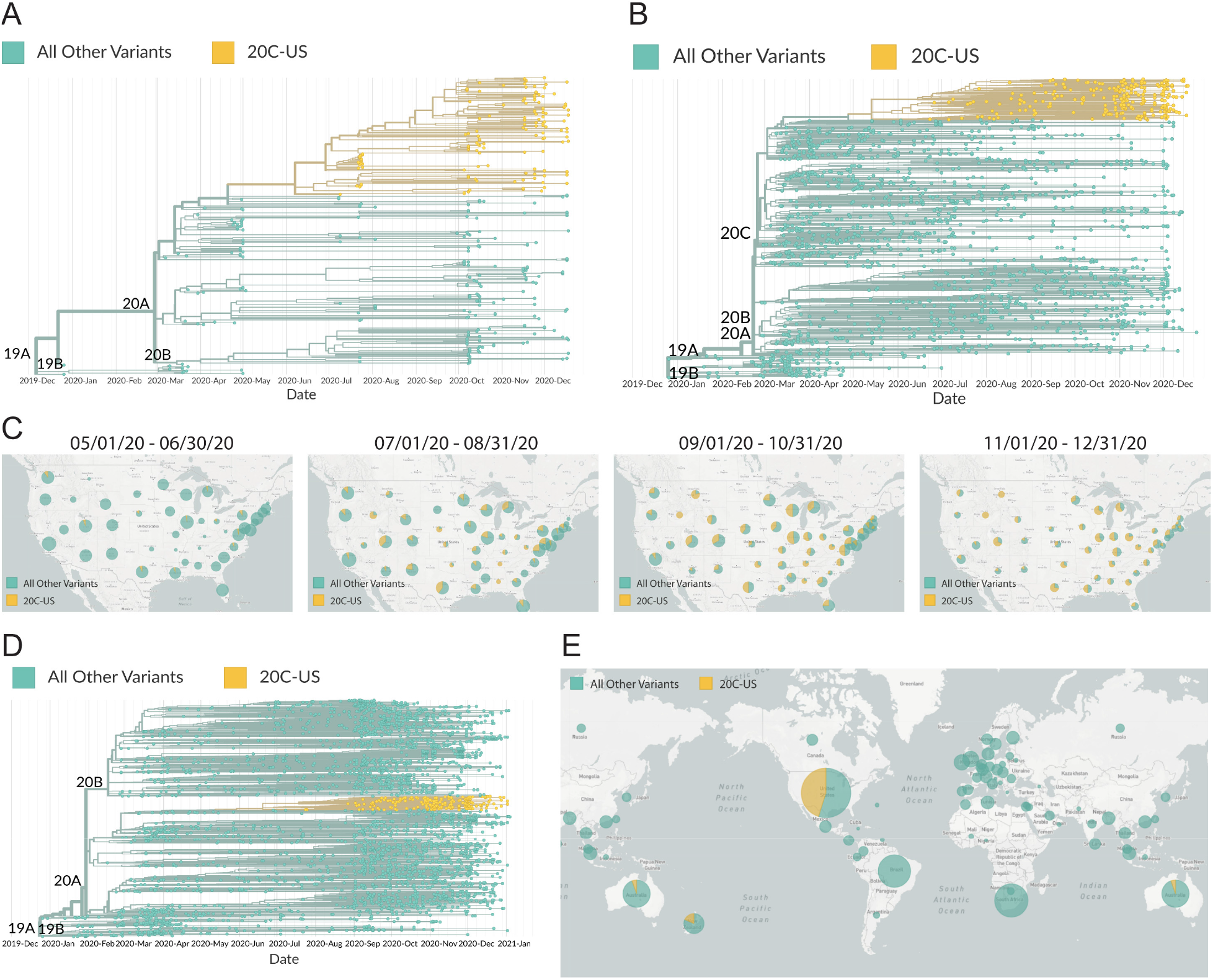
Phylogenetic reconstruction and geographic visualization of SARS-CoV-2 variant 20C-US in Illinois, the United States, and globally. Phylogenetic reconstruction of SARS-CoV2 using (**A**) 350 genomes sequenced from Illinois from March 2020 through December 2020 and (**B**) genome sequences randomly subsampled from the U.S. at ~3.3% of all U.S. genomes in the GISAID database (as of Jan. 4, 2021). (**C**) Geographic visualization of the fraction of 20C-US variant genomes in the U.S. during the indicated 2-month intervals. (**D**) Phylogenetic tree reconstruction of SARS-CoV-2 during the 2-month interval of Nov. 1 to Dec. 31, 2020 using 3819 randomly subsampled global genomes from GISAID (as of Jan. 4, 2021). (**E**) Global map view of the fraction of 20C-US variant genomes from around the world. For all trees and maps, the signature mutation ORF1b:N1653D was used to highlight the 20C-US variant. substantial fraction of genomes comprised this new variant for most U.S. states (**Fig. 1C**).

**Table 1.** Signature 20C-US mutations.

| **Mutation Name** | **Nucleotide Change** | **Amino Acid Change** | **Protein Name** | **Protein Function** |
| --- | --- | --- | --- | --- |
| **ORF1b:N1653D** | A18424G | N1653D | ORF1b, nsp14 | RNA proofreading, RNA capping, methyltransferase |
| **ORF1b:R2613C** | C21304T | R2613C | ORF1b, nsp16 | RNA capping, methyltransferase |
| **ORF3a:G172V** | G25907T | G172V | ORF3a | cellular immune modulation, virus maturation and exit |
| **N:P67S** | C28472T | P67S | nucleocapsid | viral particle structure, RNA binding |
| **N:P199L** | C28869T | P199L | nucleocapsid | viral particle structure, RNA binding |
| **ORF14:Q64\*** | C28869T | STOP | ORF14 | unknown |

To better understand this new variant at the national level, we randomly subsampled approximately 3.3% of available U.S. genomes (1905 out of 57754 genomes) from the GISAID database and constructed a phylogenetic tree using the Nexstrain pipeline (*9*) (**Fig. 1B**). Analysis of each mutation within the tree and visualization of geographic distribution revealed that a substantial fraction of genomes comprised this new variant for most U.S. states (**Fig. 1C**). When a unique, signature mutation for the new variant, ORF1b:N1653D, was visualized on a globally subsampled phylogenetic tree, a subclade of the 20C branch of the B.1.2 lineage corresponding to the new variant became apparent (**Fig. 1D**). Geographic visualization from Nov. 1 to Dec. 31, 2020 revealed that nearly half of all recently sequenced SARS-CoV-2 genomes from the U.S. are the new variant of interest (**Fig. 1E**). Furthermore, this variant has thus far only been reported at very low levels in a handful of other countries, including Mexico, Australia, New Zealand, Singapore, Thailand, Taiwan, Poland and Israel. Therefore, we have termed this new variant 20C-US.

### Tracing a timeline and geographic origin of the 20C-US variant

We attempted to establish a timeline and geographic origin for the emergence of 20C-US by tracing the appearance of mutations in sequenced genomes. We selected all SARS-CoV-2 genomes from the global GISAID database that possessed the five hallmark mutations of 20C-US as well as two prerequisite mutations that appear key to the 20C-US lineage based on phylogenetic reconstructions, which are ORF8:S24L and ORF1a:L3352F. The majority of genomes bearing these two mutations were from Minnesota and Louisiana during March and April, 2020. The first signature mutation of 20C-US, ORF3a:G172V, then appears in four genomes from Louisiana and one from Arizona in April. A series of genome sequences are then reported in late May and early June that contain the remaining four signature mutations simultaneously, suggesting either a rapid succession of mutations or an event where most or all were acquired in a single patient. Those genomes primarily originated from Texas samples, with the earliest being from the greater Houston, Texas area on May 20, 2020. The new 20C-US variant then becomes prevalent in SARS-CoV-2 genome sequences across the U.S. over time (**Fig. 1C**).

### Identification of recent mutations of potential interest in the 20C-US phylogenetic lineage

To further characterize 20C-US, we identified all GISAID samples with the signature mutations of ORF3a:G172V, ORF1b:N1653D, and N:P67S and that also possessed N:P199L or any mutations at position 2613 for ORF1b. We then reconstructed a phylogenetic tree with these 4681 sequences. A branching event was observed in this new tree that was initiated by two new mutations, a synonymous mutation at the nucleotide level, C14805T, that co-occurs at the same time with a non-synonymous mutation of the ORF1a gene that changes M2606 to I2606 (**Table 2**, **Fig. 2A**). The co-occurrence of these two mutations in the 20C-US lineage was first observed in late June of 2020. The first full 20C-US genome reported with this mutation and possessing the expected genotype was from Wisconsin on June 23, 2020 followed by an Illinois case on June 25, 2020. Visualizing the geographic distribution of 20C-US genomes with the ORF1a:M2606I mutation demonstrated that it has achieved high prevalence in the eastern and midwestern regions but has not yet spread widely to the western and southwestern regions of the U.S. (**Fig. 2B**). The ORF1a:M2606I mutation accounts for 48% of all 20C-US genomes.

**Fig. 2.**
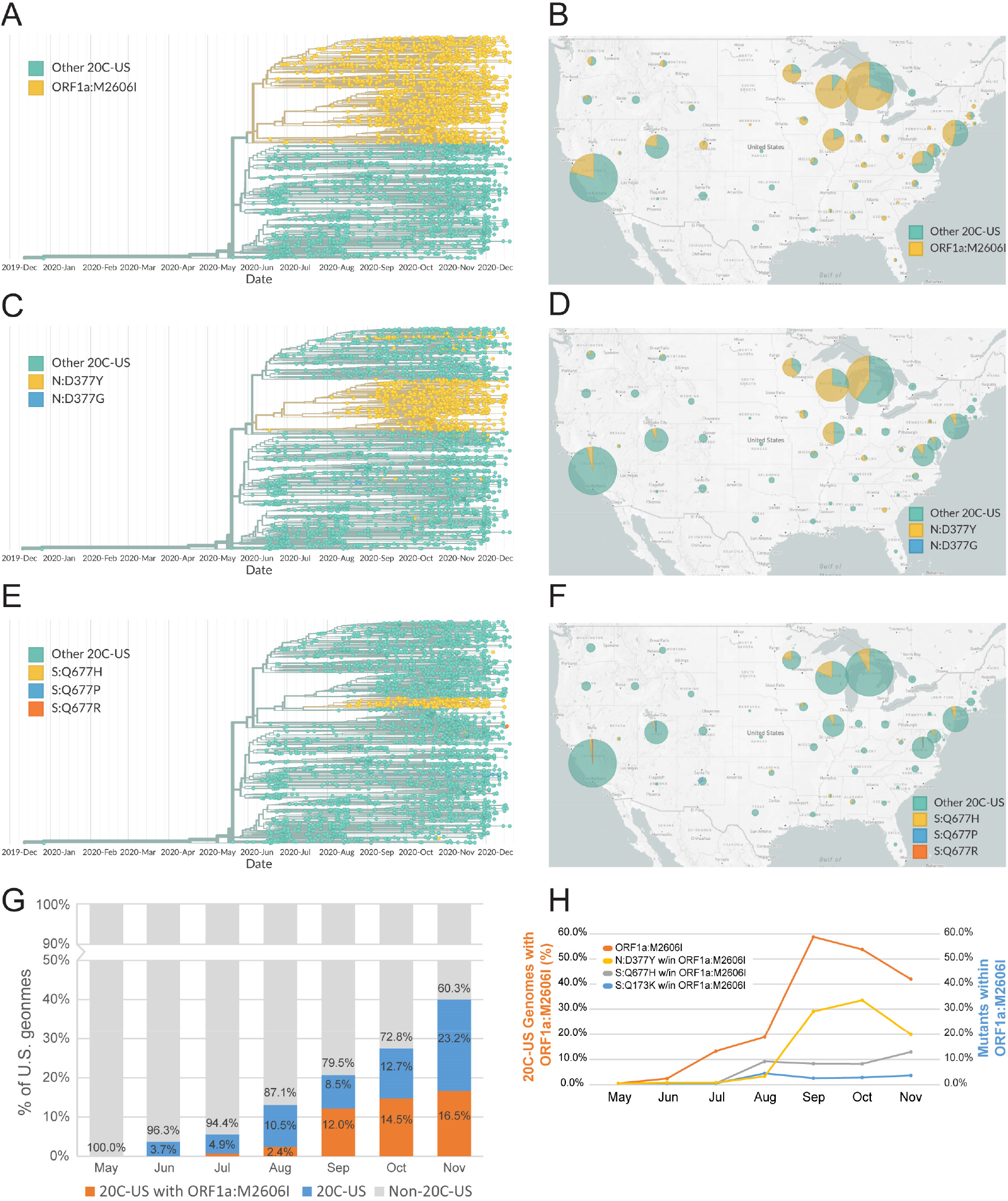
Characterization of recently acquired mutations of the SARS-CoV-2 variant 20C-US. (**A**-**F**) Phylogenetic reconstruction and geographic visualization (from Nov. 1 to Dec. 31, 2020) of all SARS-CoV-2 variant 20C-US genomes (n=4683) in the GISAID database (as of Jan. 4, 2021). The ORF1a:M2606I, N:D377Y, and S:Q677H mutant genotypes are colored to distinguish them from all other genetic variants within the 20C-US tree. (**G**) Plot depicting the rise in percentage of genomes for 20C-US and 20C-US possessing ORF1a:M2606I for the entire U.S. during the indicated months (as of Jan. 4, 2021). (**H**) Percentage of 20C-US genomes that possess the ORF1a:M2606I or N:D377Y mutation within 20C-US, as well as the the percentage of ORF1a:M2606I mutants that also possess the S:Q677H or S:Q173K mutation, versus time.

**Table 2.**
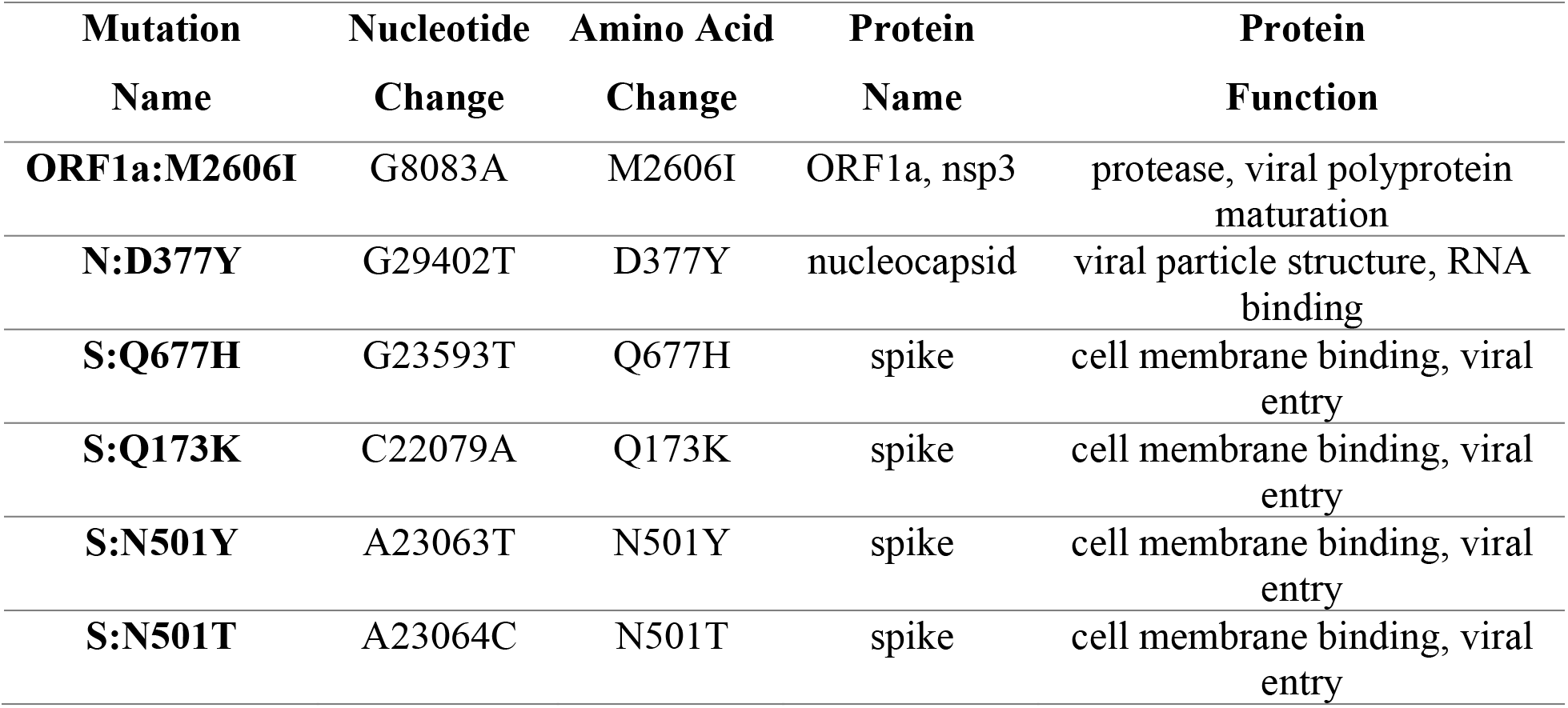
Additional new 20C-US mutations of interest.

| **Mutation**<br>**Name** | **Nucleotide**<br>**Change** | **Amino Acid**<br>**Change** | **Protein**<br>**Name** | **Protein**<br>**Function** |
| --- | --- | --- | --- | --- |
| **ORF1a:M2606I** | G8083A | M2606I | ORF1a, nsp3 | protease, viral polyprotein maturation |
| **N:D377Y** | G29402T | D377Y | nucleocapsid | viral particle structure, RNA binding |
| **S:Q677H** | G23593T | Q677H | spike | cell membrane binding, viral entry |
| **S:Q173K** | C22079A | Q173K | spike | cell membrane binding, viral entry |
| **S:N501Y** | A23063T | N501Y | spike | cell membrane binding, viral entry |
| **S:N501T** | A23064C | N501T | spike | cell membrane binding, viral entry |

Within the new clade defined by ORF1a:M2606I, three additional branches of significance emerge. One is a nucleocapsid mutation at position 377, converting an aspartate (D) to tyrosine (Y) (**Table 2**). Since the summer months of 2020, this mutation has occurred many times in the 20A, 20B, and 20C clades. However, it has clearly established a distinct and well-developed branch within the ORF1a:M2606I lineage of 20C-US (**Fig. 2C**). When viewed geographically, its distribution closely mirrors that of the ORF1a:M2606I 20C-US genotype (**Fig. 2D**).

Two other mutations that have established more recent branches within the same lineage as ORF1a:M2606I are Q677H (**Fig. 2E**) and Q173K mutations in the spike (S) gene (**Table 2**). The earliest 20C-US genomes in the GISAID database having ORF1a:M2606I and S:Q677H originated from Minnesota and Wisconsin on August 17, 2020. Mutation of Q677 to lysine (K), arginine (R), proline (P), or histidine (H) has occurred spontaneously throughout the pandemic in many lineages, including branches of the 20A, 20B, and 20C clades. However, an S:Q677 mutation has never established and expanded well until recently in the ORF1a:M2606I lineage. Within the ORF1a:M2606I lineage, the S:Q677H mutation represents 10.2% of genomes and a map view illustrates that S:Q677H mutants remain largely localized to the upper Midwest, which includes Minnesota, Wisconsin, and Michigan (**Fig. 2F**). Both N:D377Y and S:Q677H have appeared many times but only established as strong branches after 20C-US acquired the ORF1a:M2606I mutation, suggesting the possibility of an epistatic interaction between these mutations (*10*). The S:Q173K mutation is rather unique for SARS-CoV-2 and also falls in the ORF1a:M2606I lineage of 20C-US. It has established a small but growing branch in the 20C-US clade. Finally, 20C-US has also begun harboring a number of recent mutations to the S gene outside of the ORF1a:M2606I lineage that convert asparagine (N) 501 to either tyrosine (Y) or threonine (T). Thus far, fifteen recent 20C-US genomes have been found with S:N501Y/T mutations, several with S:N501Y from Texas on the same branch of the 20C-US phylogeny.

To better understand the emergence of new mutations, we plotted the percentage of 20C-US genomes possessing ORF1a:M2606I versus time for the U.S. (**Fig. 2G-H**). Interestingly, we observed an apparent slowing or reduction in the percentage of 20C-US variants carrying the ORF1a:M2606I mutation. Performing a similar analysis for N:D377Y revealed a similar trend. If decreases in the fraction of a variant over time are taken as an indication of potential fitness loss (*11*), then this mutation may not be improving the fitness of its parent lineage. However, the same analysis for S:Q677H and S:Q173K (as a percentage of US-20C genomes that also carry the ORF1a:M2606I mutation) revealed that these two mutations are not decreasing at a similar rate as the ORF1a:M2606I and N:D377Y mutants of 20C at this time (**Fig. 2H**), suggesting a potential increase in fitness. It is possible that genetic drift or a lag in genome sequence reporting could account for these aberrations, especially when so few genomes containing S:Q677H and S:Q173K in the 20C-US lineage are currently available. Indeed, both could track more closely with ORF1a:M2606I as more genome sequences become available. However, it is also possible that ORF1a:M2606I results in slightly lower fitness compared to its parent 20C-US lineage over time and the more recent spike protein mutations may compensate to help rescue fitness. Better resolution of these data through additional sequencing over time should determine the fate and impact of these novel mutations.

### Biological implications of 20C-US mutations

Several mutations carried by the 20C-US variant suggest biochemical or functional impacts on SARS-CoV-2 biology. ORF1b:N1653D is highly unique and specific to the 20C-US lineage. This mutation occurs at residue 138 in the ExoN domain of nonstructural protein (nsp) 14, a novel RNA proofreading domain that ensures the integrity of RNA genomes and transcripts (*12*). ExoN inactivation is lethal to SARS-CoV-2 (*13*). The mutation of asparagine (N) to aspartate (D) at this position represents conversion from a neutral to a negatively charged residue in a low complexity domain, possibly involved in mediating protein-protein or RNA-protein interactions (*14*) (**Fig. 3A**). In addition, nsp14 plays a critical enzymatic role by installing a methyl group on the base of the 5’ guanine cap for viral RNA transcripts, which is essential for translation into viral proteins (*12, 13*). Interestingly, the other essential viral factor involved in cap formation on viral transcripts is nsp16, which catalyzes 2’-*O*-methylation of the nucleotide adjacent to the terminal 7-methyl-guanine cap on viral RNA transcripts (*12, 15*). The signature 20C-US mutation ORF1b:R2613C results in a conversion of arginine (R) to cysteine (C) at residue 216 of nsp16, which would be predicted to disrupt hydrogen bonding to an adjacent glutamate and structured water molecule and possibly alter local protein stability (**Fig. 3B**). These two rather unique mutations co-occur in 20C-US and could potentially alter genome integrity, mutation retention, transcript integrity, and translation efficiency of viral messenger RNA.

**Fig. 3.**
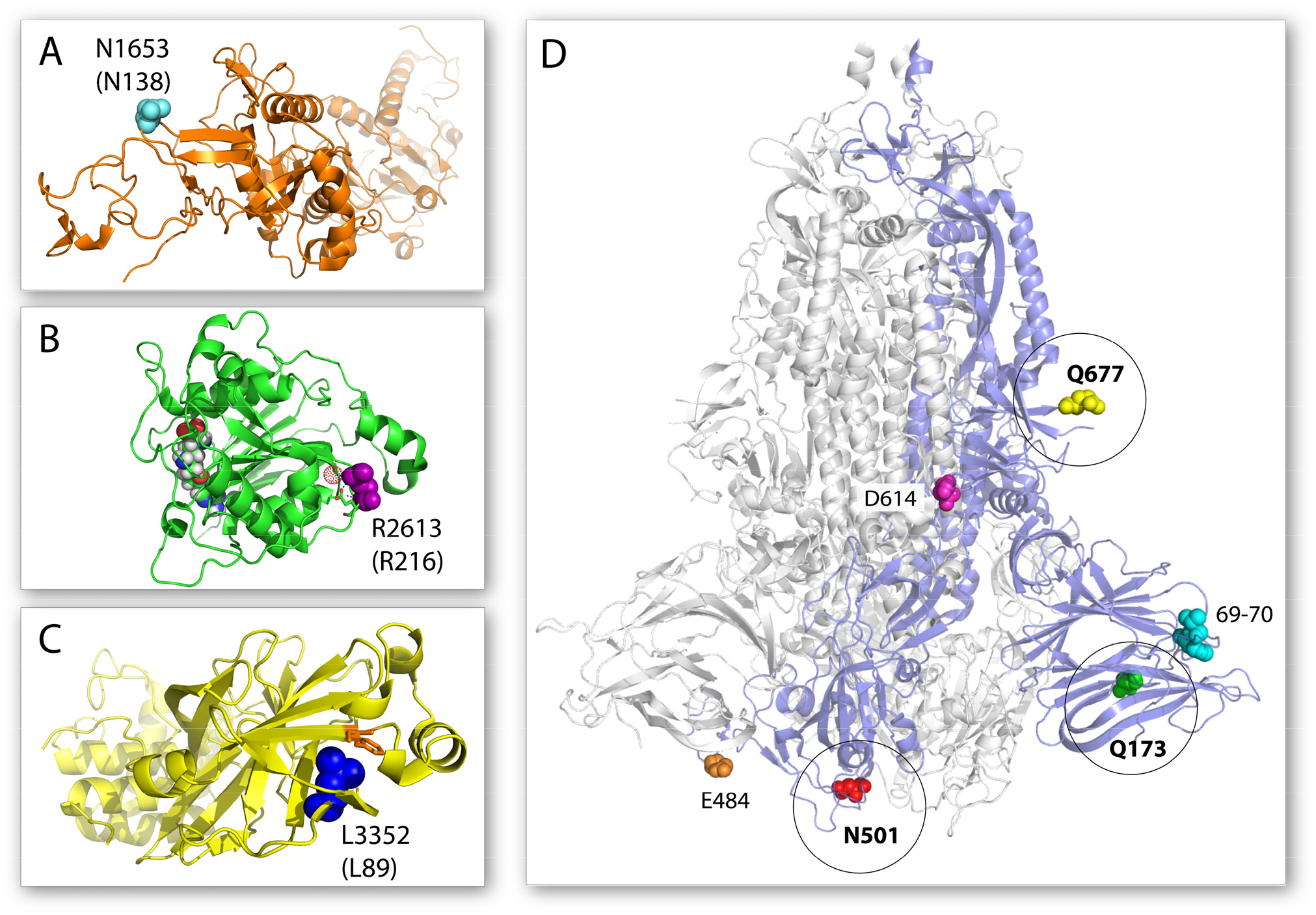
Location of select 20C-US mutations on structures of their respective proteins. (**A**) Structure of nsp14 and the N138 amino acid shown in cyan spheres (PDB 5C8S). (**B**) Structure of nsp16 and the R216 amino acid shown in purple spheres (PDB 6YZ1). The neighboring glutamate residue and the structured water molecule that it is predicted to hydrogen bond with are shown as green sticks and a red dot sphere, respectively. An S-adenosyl-methionine analog is shown as a space filling model in the nsp16 active site. (**C**) The structure of nsp5 and the L89 amino acid shown in blue spheres (PDB 7KHP). A nearby phenylalanine available for increased hydrophobic packing is shown as orange sticks. (**D**) Structure of the trimeric spike protein with one monomer shown as a blue cartoon (PDB 7JJI). The positions of the Q677, Q173, and N501 amino acids are shown in yellow, green, and red spheres, respectively. Additional important mutations of the spike protein described previously are also shown: the 69-70 amino acid deletion in cyan spheres, the E484 amino acid in orange spheres, and the D614 amino acid in magenta spheres.

The two largest SARS-CoV-2 viral RNA transcripts, ORF1a and ORF1b, are translated into polyproteins that must be further processed by proteases to release mature, functional viral nsp proteins. Two proteases within the ORF1a gene are responsible for this processing, nsp3 and nsp5 (*12*). The parental ORF1a:L3352F mutation carried by 20C-US creates a mutation of leucine (L) to phenylalanine (F) at residue 89. L89 in nsp5 is involved in hydrophobic packing of a domain adjacent to the expected active site. There appears to be room to accommodate a larger phenyl ring. Packing against another juxtaposed F side group in the domain could conceivably improve hydrophobic packing and enhance protein stability (*16*) (**Fig. 3C**). The more recently acquired mutation of ORF1a:M2606I results in a mutation to the rather large nsp3 protein within what is predicted to be the C-terminal 3Ecto or Y domain. This domain is implicated in anchoring the replication-transcription complex (RTC) to the endoplasmic reticulum membrane to facilitate interaction of nsp3 with other viral proteins on the cytosolic side (*17*).

ORF3a is a multifunctional protein involved in several aspects of the viral life cycle at the surface of the cell membrane and intracellular membranes (*18*). ORF3a modulates the innate immune response and apoptosis of host cells (*19*). The mutation G172V in ORF3a occurs within a conserved di-acidic Asp-X-Glu domain (171-173) involved in trafficking to the cell membrane (*20, 21*). The introduction of a valine (V) at the more variable 172 position may modulate interactions with viral or cellular factors (*22*). ORF3a plays a role in viral particle maturation and release at the cell membrane and has been proposed to co-mutate with the spike protein (*23*). Substitution of serine to leucine at position 171 of ORF3a is a common mutation of the South African SARS-CoV-2 variant 501Y.V2.

The 20C-US variant has also recently acquired a Q677H or a Q173K mutation in the spike protein. Q677H is directly adjacent to the furin cleavage site. The furin cleavage site is a novel motif not observed in SARS-CoV viruses that is proposed to significantly enhance infectivity (*24*). Furin cleavage is a critical priming step essential for efficient entry of SARS-CoV-2 viruses into cells (*24*). A mutation of interest has been the P681H in the spike protein of the novel UK variant 501Y.V1, also due to close proximity to the furin cleavage site (*5*). Q677 and P681 are mutated to histidine in 20C-US and 501Y.V1, respectively, suggesting a potentially important effect of histidine near the furin cleavage site. The Q677 amino acid resides in a similar region on the spike protein as D614, which is commonly mutated to a G residue (*3*) (**Fig. 3D**).

S:Q173K resides in the S1^A^ domain nearby the receptor binding domain (RBD), where the well-known E484K and N501Y mutations are found (*5, 6*). The S1A domain, like the RBD, exhibits low conservation, which helps SARS-CoV-2 adapt to host cells and host immunity. Although antibodies against the SARS-CoV-2 spike protein are believed to primarily target the RBD (*25*), antibodies isolated from COVID-19 convalescent patients have been found to bind very tightly to the S1^A^ domain (*26*). Conversion from a neutral to charged amino acid might alter interactions important for antibody recognition. In addition, N501Y and N501T mutations in the spike protein are beginning to occur in the 20C-US variant. The significance of S:N501 mutations has been underscored by their reoccurrence in apparently highly transmissible forms of the SARS-CoV-2 virus (*5, 6*). These mutations occur in the RBD and are directly implicated in modulation of host cell interaction (**Fig. 3D**).

## Discussion

We have characterized the emergence and rise of a prevalent SARS-CoV-2 variant within the 20C clade that is highly specific to the continental U.S. 20C-US is predicted to soon surpass 50% penetrance to become the dominant variant in the U.S. (**Fig. 4A**). It is unclear whether natural selection or genetic drift has driven the rise in prevalence of 20C-US. Nonetheless, its dominance has been largely achieved during the third pandemic wave when cases have risen significantly (**Fig. 4B**). During this period, Google mobility data are consistent with no major changes in population movement patterns across the U.S. that could account for an increase in the proportion of 20C-US to other variants (**Fig. 4C**). Recent studies on hospital care for COVID-19 patients in the U.S. indicate that adjusted mortality rates are decreasing and patient outcomes are improving (*27, 28*). Taken together, these observations suggest the possibility that 20C-US may have some degree of increased transmissibility but not a significantly increased disease severity.

**Fig. 4.**
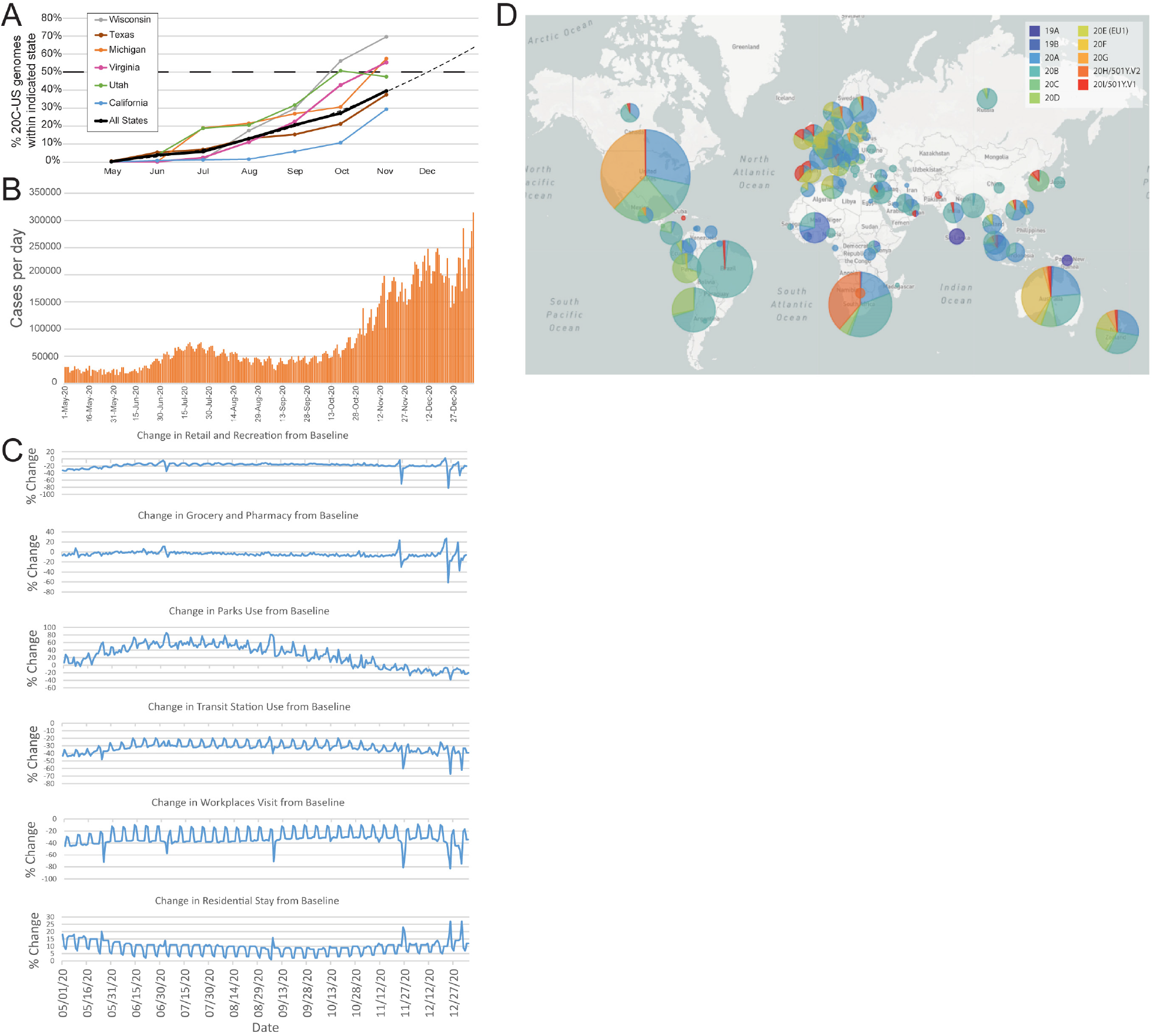
20C-US rise in prevalence correlated with population mobility, disease severity approximations, and global variant emergence. (**A**) Plot depicting the rise in percentage of 20C-US genomes for all SARS-CoV-2 genomes from the U.S. in the GISAID database in the indicated states and indicated months up to Nov. 30, 2020. A curve fit to the average of all U.S. states projecting the continued rise of 20C-US genomes reported in the U.S. (**B**) Plot of confirmed daily cases of COVID-19 in the U.S. (https://covid.cdc.gov/covid-data-tracker/#trends_dailytrendscases). (**C**) Google mobility plots for diverse activities for the entire U.S. Data is presented as percent change from pre-pandemic baseline. (**D**) Global map view of using Nextstrain’s designated SARS-CoV-2 clades from Aug. 1 to Dec. 31, 2020 using a phylogenetic tree reconstruction (n=3819 randomly subsampled genomes in the GISAID database).

The biological effects of the combined 20C-US mutations, as well as the viral characteristics of 20C-US, like fitness, transmissibility, and virulence, remain to be experimentally characterized. However, we note that all signature mutations that occurred during the establishment of the 20C-US variant lineage were non-synonymous. These include mutations to proteins involved in viral replication, metabolism, and cellular exit. Importantly, one mutation, ORF1b:N1653D, occurs in the RNA genome proofreading enzyme nsp14. 20C-US has more recently also acquired additional mutations. Of particular interest are independently arising Q677H, Q173K, and N501Y/T mutations in the spike protein. Q677H lies near the furin cleavage site that could potentially alter cellular entry of virus particles. Q173K and N501Y/T reside within the N-terminal S1^A^ and RBD domains, respectively, and have the potential to alter antibody recognition. Monitoring the fate of these mutations, as well as the potentially linked M2606I mutation of ORF1a, within 20C-US should provide critical insight into ongoing evolutionary processes for SARS-CoV-2 and their impact on real-world pandemic outcomes.

Although characterization of distinct novel SARS-CoV-2 variants has been limited up to this point, two driving forces that may affect their rise and prevalence are the near simultaneous acquisition of multiple mutations and limited population movement between local regions during the pandemic. The UK variant 501Y.V1 acquired several mutations very rapidly. It has been proposed that this founding event occurred in an immune-compromised patient where sufficient pressure to continually evade the immune system was present (*4*). The South African variant 501Y.V2 also quickly acquired multiple mutations in the spike protein (*6*). Likewise, the 20C-US variant appears to have incorporated five mutations very rapidly. These events might create a jump in fitness and a temporary disequilibrium in variant competition.

The mechanism and rate of SARS-CoV-2 transmission necessitates strict measures that effectively limit population movement (*29*). Since the outbreak of the global COVID-19 pandemic, international travel has become highly restricted. Novel variants that emerge in an isolated region or country may transmit locally among that population and develop distinct genotypes and phenotypes. Thus, it would be expected that regional territories would develop their own distinct SARS-CoV-2 variants over time. When searching for novel emerging variants, focusing on local and regional data may provide an advantage. Our ability to identify the 20C-US variant can be partly attributed to our initial focus on Illinois state-level data since the prevalence of the 20C-US variant was more pronounced in the U.S. Midwest.

While this manuscript was in preparation, the Nextstrain group updated their global phylogenetic analysis server for SARS-CoV-2 to begin designating emerging clades. We found that the 20C-US variant closely tracks with the newly designated 20G clade, demonstrating that this approach will be valuable in helping to identify new potential variants or clades of interest. When visualized on a global geographic view, it becomes clear that variants specific to other world regions have slowly emerged and gained prevalence, including the western coast of South America (20D), Europe (20E/EU1), Australia (20F), South Africa (20H/501Y.V2), and the United Kingdom (20I/501Y.V1) (**Fig. 4D**). A detailed assessment of the emergence and rise to prevalence should also be undertaken for these variants. Unless successful vaccination efforts can be greatly accelerated, we predict the emergence of dominant novel variants in many global regions that are relatively isolated, possibly including Brazil, New Zealand, the African west coast, and Japan.

This study underscores the need for greater genomic surveillance of the SARS-CoV-2 virus, especially at the regional level where novel variants will first emerge. Modern genomic surveillance enables observation of evolution in near real-time, prediction of major shifts in viral fitness, and assurance that vaccines are kept current.

## Materials and Methods

### Sequencing of SARS-CoV-2 Samples

RNA extraction was performed using Mag-Bind Viral RNA XPress Kit (Omega Bio-Tek) on samples received from Illinois Department of Public Health (IDPH) in inactivation buffer consisting of 200 μL of sample in Viral Transport Media (VTM) and 240 μL of lysis buffer (239 μL of TNA lysis buffer and 1 μL of carrier RNA per sample). cDNA synthesis from the extracted RNA was performed using ABI High-Capacity cDNA Reverse Transcription Kit, following manufacturer’s recommended protocol. SARS-CoV-2 was detected using N2 primers (Integrated DNA Technologies, IDT) that target the N2 region of the nucleocapsid gene. PrimeTime Gene Expression Master Mix (IDT) and 2 μL of cDNA was used to determine the *Ct* value of each sample. Sequencing was performed with Oxford Nanopore Technology’s MinION platform using ARTIC Network protocol (https://artic.network/ncov-2019) with slight modifications to the protocol. Briefly, 25 μL reactions were performed for each pool using 5 μL of cDNA template and 4 μL of 10 μM primer pool in respective reactions. Finally, 0.5 μL of 10 mM Deoxynucleotide (dNTP) solution mix (New England Biolabs) was added to each reaction and nuclease -free water was used to make up the remaining reaction volume. The PCR was run by combining the annealing and extension steps at 63°C. The thermocycler was set at 98°C for initial denaturation for 30 seconds, followed by 35 cycles of denaturation at 98°C for 15 seconds, annealing and extension at 63°C for 5 min and holding at 4°C indefinitely. From each Multiplex PCR, 5 μL of several reactions from each pool from the batch was used to run on 1% agarose gel. The remaining 20 μL of each pool were pooled and clean-up was performed using an equal volume of AMPure XP beads (Beckman Coulter) and quantified using 1 μL of sample with the Qubit dsDNA HS Assay Kit on a Qubit 2.0 Fluorometer. Following quantification, 60 ng of each sample was end-prepped using 0.75 μL of Ultra II End-Prep enzyme and 1.75 μL of Ultra II End-Prep buffer with a total volume of 15 μL. The samples were then barcoded using the 96 native barcoding kit (Oxford Nanopore Technology) and further processed using the ARTIC Network protocol (https://artic.network/ncov-2019).

Basecalling on completed sequencing runs was performed using Guppy high accuracy model and further demultiplexed using Guppy barcoder using strict parameters requiring barcodes to be present at both ends. Reads were filtered based on length quality (400-700 bp) and mapped by aligning to the MN908947.3 reference genome using minimap2. Medaka was used to create the consensus sequence and call variants for each of the samples. The variants identified were fed into longshot to produce a set of high-confidence variants.

### Dataset

SARS-CoV-2 genomes sequences used in this paper that were not generated by us were acquired from the GISAID Initiative (https://www.gisaid.org/). This dataset was updated on 2021-01-04. Individual sequences were compiled by GISAID from contributions from individual labs across the world. Their information is compiled in the supplementary section. Sequences generated by our laboratory, currently in the process of being submitted to GISAID, were added to this dataset. Our samples were all within the state of Illinois.

All sequences were evaluated through the command-line version of Nextstrain’s NextClade software (https://www.npmjs.com/package/@neherlab/nextclade) (*9*) to derive a list of all nucleotide and amino-acid substitutions. 20C-US samples were then filtered from the larger population using a criterion of necessary amino-acid mutations. Because not all sequences are complete (contain gaps) there is the possibility of a sequence being within the 20C-US population but not reporting one of the required mutations. The formula for filtering therefore used a flexible criterion.

### Phylogenetic Trees and Maps

Once appropriately filtered from the greater genomic sample pool, sample FASTA files were used for phylogenetic inference. Briefly, the Augur pipeline utilized by Nextstrain filters sequences for metadata values and N content before aligning them using MAFFT (*30*). As a reference, the Nextstrain nCoV toolkit aligns SARS-CoV-2 sequences to the SARS-CoV-2 isolate Wuhan-Hu-1 complete genome (GenBank: NC_045512.2) and by default removes any nucleotide insertions relative to this sequence. The pipeline conducts a maximum likelihood phylogenetic analysis using the general time reversible model allowing for invariant sites and a gamma distribution (GTR+I+G) in IQ-TREE (*31*). Augur further refines and annotates the tree in various ways. The resulting files are combined and used by Auspice to visualize phylogenetic relationships and geographic distributions of SARS-CoV-2 across time. For the 20C-US subset, all samples were used. For the model of all SARS-CoV-2 variants within the United States of America, a subset of the data was generated based on Nextstrain’s standard filtering criteria. Briefly, lower-quality samples (those with large numbers of gaps or lacking sufficient metadata) are filtered out, then the dataset is sorted based on the month it was acquired as well as the U.S. state it was acquired in. From each of these subsets, Nextstrain attempts to randomly pick an equal number of samples. All these samples are then recombined and processed together.

## Supporting information

GISAID author credits

## Acknowledgments

We thank the Illinois Department of Public Health for access to patient samples.

## Author contributions

A.A.P planned and performed aspects of all experiments and genomic analyses, prepared figures, and assisted in writing the manuscript. M.S.B. analyzed sequencing data, analyzed GISAID and local datasets, prepared figures, and assisted with manuscript writing. C.L.B. performed sequencing runs, analyzed phylogenetic and genomic data, and assisted in writing of the manuscript. K.N.O. designed and performed early experiments in the sequencing workflow, including RNA extractions, PCR and qPCR. R.C., M.P., S.B.E., A.O.Y., A.A.W., P.E.M., R.J.S., M.M.H., and S.W.B. all performed one or more individual molecular biology steps in the genome sequencing workflow. K.S. accessed and analyzed Google mobility and COVID-19 case data. K.T.G. conceived, planned, and designed experiments, as well as performed experiments, prepared figures and wrote the manuscript.

## Data and materials availability

Genome sequences are currently being submitted to, and will be available through, the GISAID initiative (https://www.gisaid.org/). Gagnon laboratory Illinois genome sequence results can be visualized and analyzed on their Nextstrain group page: https://nextstrain.org/groups/illinois-gagnon-public.

